# DIGEST: An online tool for designing of multiple reaction monitoring assays

**DOI:** 10.1101/2023.11.27.568790

**Authors:** Prutyay Gautam, Praveen Singh, Mrinal, Akash Bhaskar, Sukriti Sacher, Yashesh Dagar, Trayambak Basak, Shantanu Sengupta, Arjun Ray

## Abstract

Targeted proteomics using multiple reaction monitoring (MRM) assays enables fast and sensitive detection of a preselected set of target peptides. This technique utilizes the specificity of precursors to product transitions for quantitative analysis of multiple proteins in a single sample. The success of an MRM experiment depends on the selection of transitions however, given the existing resources, accurately predicting signal intensity of peptides and their fragmentation patterns ab initio is challenging task. We present an alternative for rapid design of MRM transitions for proteomics research: DIGEST. Our method predicts the b and y ions with +1 and +2 charge produced in a collision cell of a mass spectrometer from peptides of multiple proteotypically digested proteins. Additionally, by using the existing knowledge of the fundamental rules for designing transitions, the tool provides optimal MRM transitions, negating the need to undertake prior “discovery” MS studies. We demonstrate that our algorithm is directed toward the selection of MRM precursor and product-ions pairs, and can avoid the pitfalls of interference due to cross-contamination of samples by selecting ion combinations that uniquely map to target peptides. Comparison with SRMAtlas showed that DIGEST successfully predicted the peptide and production pairs in the majority of cases. We believe that DIGEST will facilitate rapid design of MRM assays with increased specificity, reducing the overall time required to design an MRM assay for routine mass-spectrometry. DIGEST is available as a web-based tool at https://digest.raylab.iiitd.edu.in/

## INTRODUCTION

Recent developments in mass spectrometry (MS)-based proteomics have enabled systems biology approaches and biomarker discovery applications^1^. However, identifying potential biomarker candidates with robustness, reproducibility and sufficient throughput presents several challenges^2,3^. Multiple reaction monitoring (MRM) assays have emerged as a powerful technique due to their high-throughput multiplexing capabilities for quantification of several proteins in a single liquid chromatography mass spectrometry (LC-MS) run^3^. MRM is usually performed on triple-quadrupole or quadrupole-ion trap mass spectrometers and uses the strategy for targeted data acquisition using transitions (filtering precursor ion and fragment ion pair)^4^. The focused protocol for targeted (MRM based) proteomics approaches, starts with selection of a list of transitions (Q1/Q3), in contrast to acquisition of a full fragment ion spectrum, leading to a reduction in assay time and redundancies in data collection with increased sensitivity and reproducibility across multiple runs. MRM has an edge over traditional immunoaffinity-based techniques for validation of biomarkers. Immunoaffinity assays struggle with issues such as cross-reactivity, complexity in multiplexing and a lower dynamic range of proteins in the screening panel. In contrast, MRM can quantify multiple proteins that are less abundant, their isoforms, and post-translational modifications accurately in a single run^3,4^. Therefore, targeted proteomics utilizing MRM assays hold immense potential in the transition of proteomics from a discovery-oriented technique to a robust and quantitative method suitable for hypothesis-driven studies in contemporary biological research.

One of the major requirements of an MRM based-targeted proteomics workflow is the compilation of the list of target proteins and peptides, and the necessary attributes of these targets to facilitate the measurements. Additionally, MS platform variability in reliably and consistently detecting unique peptides for the identification of proteins adds to this problem^5^. Previously, it has been well demonstrated that unique ion signatures (UIS) representing a combination of ions that map exclusively to one peptide can be used to quantify the target protein. Since MRM assays measure these unique transitions, a significant amount of time is spent in designing and measuring transitions for specific peptides that are both selective and sensitive. Historically, these precursor-product pairs have been predicted using heuristic approaches followed by a refinement after the initial discovery phase experiment^6^. Moreover, if only a few transitions are used, it is likely that more than one peptide from a different protein may map to the selected transitions for target protein in a given MRM assay^5^. Moreover, since triple-quadrupole instruments work with limited resolution, peptides that have similar transitions close to Q1 and Q3 values being used to identify the target peptide, may lead to ambiguity in detection. Therefore, MRM transitions for a peptide should be carefully chosen with minimal redundancy and interference with other peptides^7^.

Utilization of in-silico approaches for selecting transitions for a successful MRM experiment has shown promise^8–11^. Among these TIQAM (Targeted Identification for Quantitative Analysis by MRM) software suites support MRM workflow by generating transitions from in-silico digestion, by connecting to PeptideAtlas for transition selection^11^. However, it uses complete MS/MS fragment spectra data which is not provided by a triple-quadrupole instrument, even though the precursor fragmentation in the collision cell is complete. Similarly, MIDAS (MRM-initiated detection and sequencing) combines an MRM scan with a full MS/MS product ion scan that can examine all fragment peptides and their corresponding transitions for sequence confirmation^12^; however, it is only compatible with a QTRAP (Quadrupole ion trap) system. Moreover, it can not be performed on triple-quadrupole systems, which are frequently used for this purpose. Popular softwares like Skyline also integrates the MIDAS or SRMcollider pipeline for targeted proteomic analysis and therefore faces the same challenges as those tools^13^. ATAQS (Automated and Targeted analysis with Quantitative SRM) provides a number of modules to support MRM assay development, however its installation and usage requires prior comprehensive computational expertise^8^. Moreover, all of these softwares are single-user desktop applications and therefore require installation on a computer. While MRMaid and SRMcollider are two web-based tools that were developed to aid in the development of MRM transitions^9,10^, however, these tools are no longer functional.

Considering there are limited options for tools that enable MRM design support without involving acquisition of MS/MS data for prediction process and that are freely accessible, we have developed DIGEST: a web-based tool that offers rapid design of MRM transitions for proteomics research. Digest can predict all the fragments b- and y-ions that can be theoretically produced in a collision cell of a mass spectrometer; from peptides of multiple proteolytically digested proteins with their frequently occurring post translational modifications (PTMs). Additionally, the program combines the knowledge of fundamental rules for designing optimal MRM transitions to provide reliable, non-redundant UIS for the peptide of interest for a given protein. DIGEST can also be used to find interfering transitions from a non-target peptide in the same proteome or for cross-contamination (from other organisms). Our tool does not need to be set up locally and does not require any computer expertise for its usage. Moreover, it can support diverse MS instruments. Using Apolipoproteins as target, we demonstrate that our algorithm carefully selects MRM precursor and product-ions pairs, and can avoid the pitfalls of interference. Furthermore, we compare the transitions obtained by DIGEST with those obtained from SRMAtlas^14^ to validate our tool’s results. We believe that our tool will be useful in the rapid designing of MRM assays that are specific. DIGEST can be accessed at https://digest.raylab.iiitd.edu.in/

## EXPERIMENTAL PROCEDURES

### Computational method

DIGEST consists of an internal database of 22 different proteomes which have been digested by 8 enzymes using Rapid Peptides Generator^15^. The proteolytic digestion by each enzyme generates theoretical precursor ions, which are stored and sorted according to their precursor ions (*m/z* values). This includes all the peptide precursor ions with zero, one and two miscleavages. For each precursor ion, fragment ions (including b and y ions with +1 and +2 charge) are generated. A *m/z* value is assigned to each of the daughter ions, fine-tuned by the user defined input of product ion (Q3) mass filters along with a tolerance level. The mass for each peptide sequence is calculated as the sum of the mono-isotopic masses of all the amino acids present in it.

All combinations consisting of three daughter ion fragments (transitions) are produced and queried with the database to identify any other precursor ions they may match with. Only those transitions are considered that have zero or single match with the database, representing unique ion signatures for the given precursor ion. All such unique ion signatures are ranked to give the optimum transition that can uniquely identify the precursor ion. If a unique transition is obtained, the user is provided with an output, or else, the size of transition combinations is incremented by one.

### Estimating MRM coverage in different organisms

To identify the minimum number of transitions needed to uniquely identify a target peptide within a proteome we performed a simulation on five different proteomes: severe acute respiratory syndrome coronavirus 2 (SARS-CoV-2), *Escherichia coli, Saccharomyces cerevisiae, Arabidopsis thaliana*, and *Homo sapiens*. Proteomes for each organism were extracted from the Swiss-Prot database^16^ and all the unique peptides post digestion for each proteome were obtained from the ExPASy Server^17^ such that their sequence length was between 6-25 amino acids, *m/z* value ranged between 300-1500, had zero missed cleavages and charge state of 2+. These post digestion peptides for each organism were subjected to the Design MRM module of DIGEST and binned according to the minimum number of transition states that were needed to identify it. The precursor ion (Q1) and product ion (Q3) tolerance value was set to 0.5 Da, while the precursor ion charge and product ion charge were chosen as 2+ and 1+ respectively.

### Experimental Methods

To validate the functionality of our tool we designed an MRM method for detection and comparative quantitation of 20 peptides from 12 human Apolipoproteins. UIS for each of the selected peptide were obtained through DIGEST using the precursor ion (Q1) and product ion (Q3) tolerance of 0.7 Da, precursor ion charge of +2 and product ion charge of +1, enzyme-Trypsin, organism-*Homo sapiens* and database-Swiss-Prot. Top UIS from the DIGEST output were incorporated in the MRM. Also, for each of the selected peptides, five most intense product-ions were selected from SRMAtlas database using ‘Complete Human SRMAtlas’ build and TSQ as targeted instrument, with the range of *m/z* value between 350-1400.

Protein from pooled plasma samples of 10 healthy individuals were precipitated using pre-chilled acetone. Briefly, to 100 µl of pooled plasma samples, 400 µl of pre-chilled acetone was added and the samples were vortex mixed and incubated at -20°C overnight. The samples were then centrifuged at 15000g for 10 minutes at 4°C. Supernatants were discarded and protein pellets were air dried at room temperature and suspended in 0.1M Tris-HCl with 8M urea and pH 8.5. Protein estimation was performed using Bradford assay. 100 µg protein was reduced with 2 mM dithiothreitol (DTT) at 56°C for 30 minutes and alkylated with 2.2 mM iodoacetamide (IAA) at room temperature in the dark for 20 minutes. The samples were diluted with 0.1 M Tris-Hcl, pH 8.5 to a final concentration of 1 M urea, and the proteins were digested overnight with sequencing grade trypsin (Promega, Madison, WI) using an enzyme: substrate ratio of 1:20. Digestion was stopped by adding formic acid to a final concentration of 0.1%. Peptide mixtures were desalted using reversed phase cartridges Oasis HLB cartridge (Waters, Milford, MA) according to the following procedure: wet cartridge with 1000 μl of 100% acetonitrile, was equilibrated with 1000 μl of 0.1% formic acid. To this acidified digest was loaded and the peptides were subsequently washed 1000 μl of 0.1% formic acid. The digested peptides were eluted 1000 μl of 70% acetonitrile in 0.1% formic acid. The peptides were dried using a vacuum centrifuge and subsequently solubilized in 100 μl of 0.1% formic acid to give a final concentration of 1 μg/μl.

Tryptic peptides were analyzed using a TSQ Altis (Thermo Fisher, San Jose, CA). The instrument was equipped with an H-ESI ion source. A spray voltage of 3.5 keV was used with a heated ion transfer tube set at a temperature of 325 °C. Chromatographic separations of peptides were performed on the Vanquish UHPLC system (Thermo Fisher, San Jose, CA). 10 μl of sample was injected in technical replicates of three and peptides were loaded on an ACQUITY UPLC BEH C18 column (130Å, 1.7 µm, 2.1 mm X 100 mm, Waters) from a cooled (4 °C) autosampler and separated with a linear gradient of water (Buffer A) and acetonitrile (Buffer B), containing 0.1% formic acid, at a flow rate of 300 µl/min for a 30 minutes gradient run.

The mass spectrometer was operated in SRM mode. For SRM acquisitions, the first quadrupole (Q1) and the third quadrupole (Q3) were operated at 0.7-unit mass resolution. A cycle time of 2 sec was chosen, and acquisitions occurred over the whole gradient of 30 min. Argon was used as the collision gas at a nominal pressure of 1.5 mTorr. Collision energies for each transition were calculated according to the following equations: CE = 0.0339 x *m/z* + 2.3398 (where CE indicates collision energy, and *m*/*z* indicates the mass to charge ratio) for doubly charged precursor ions.

Skyline was used to analyze the data. A peak was considered a true signal if it was observed in at least two of the three replicates using either ‘Design MRM’ from DIGEST or five products per precursor for SRMAtlas^14^ output data.

## RESULTS

### Minimum number of transitions required to identify a protein is three

While designing a MRM assay, deciding upon the minimum number of transitions that can form a UIS is important. In the context of optimizing the cycle time, it depends upon the number of transitions monitored and the dwell time used for scanning each precursor-product ion pair. So while monitoring a higher number of transitions or number of fragment ions per peptide may increase specificity, it may cost more time and allow detection of only a limited number of peptides in a single MRM run. As certain screening assays may require increasing the peptide numbers in a single run, intelligent ways to reduce the number of fragments in the UIS for each peptide is desirable. Previously, it has been shown that single transitions when used to identify a protein, suffer from high redundancy and lack of specificity^10^. Moreover, using combinations of two or more transitions for monitoring decreases this redundancy, increasing the specificity of the ion signature^7^. We simulated a coverage analysis to calculate the percentage of proteome that could be uniquely identified with an ion signature consisting of three, four, five or six or more than six transitions. Representative proteomes from organisms varying in complexity were chosen. **Table 1** shows that a unique ion signature consisting of three transitions could uniquely identify∼ 99% proteome of single-celled organisms (virus, bacteria and yeast) and∼97% proteome of multi-cellular organisms (*Arabidopsis thaliana*, humans). Additionally, those peptides that could not be identified with a triplet ion signature, required more than six transitions for their identification. It is possible that these peptides may be too small (<= 6 AA in length) and not proteotypic (unique to a protein) or they represent isoforms or close homologs that require a need for a complex ion signature. Therefore, we observed that a UIS made up of three transitions is sufficient to identify a protein.

**Table 1:** The percentage coverage of proteome that can be represented by a unique ion signature of three transitions.

| S.No | Organism | Total peptides | Number of peptides identified with UIS of three transitions | Number of peptides identified with UIS of more than six transitions | % of proteome covered by three transitions | % of proteome covered by more than six transitions |
| --- | --- | --- | --- | --- | --- | --- |
| 1 | SARS-CoV-2 | 486 | 486 | 0 | 100% | 0% |
| 2 | *Escherichia coli* | 29,505 | 29,462 | 43 | 99.85% | 0.14% |
| 3 | *Saccharomyces cerevisiae* | 141,330 | 139,939 | 1391 | 99% | 0.98% |
| 4 | *Arabidopsis thaliana* | 339,476 | 332,724 | 6752 | 97.95% | 1.98% |
| 5 | *Homo sapiens* | 512,994 | 500,914 | 12080 | 97.6% | 2.35% |

### Designing MRM using DIGEST

DIGEST can be used to rapidly design MRM assays with minimum inputs from the user consisting of the query peptide sequence, tolerance value for precursor ion (Q1) and product ions (Q3) and their charges. DIGEST has stored all the precursor ions generated from proteolytic digestion of a proteome with their molecular masses on binary files, with each file containing records such that the maximum difference between two entries in the file is no more than 1 Da. This makes accessing entries with the user defined Q1 input extremely efficient and fast by limiting the total number of entries to be traversed. Once the precursor and product ions, along with their *m/z* value for user defined input have been generated, transitions for product ions are generated. All combinations of product-ions consisting of three transitions are queried with the inbuilt database to identify any other precursor ion they may map to. Only those transitions are further considered that have zero or single match with the database, representing unique ion signatures for the given precursor ion. All such unique ion signatures are ranked to give the most optimum transition that can uniquely identify the precursor ion. If a unique transition is obtained in this way the user is provided with an output, or else, the size of transition combinations is incremented by one.

Therefore, DIGEST provides an ion signature consisting of either three or more transitions for the user defined input peptide with assurance that this signature is unique within the proteome. **Figure 1** summarizes the MRM design workflow of DIGEST.

**Figure 1:**
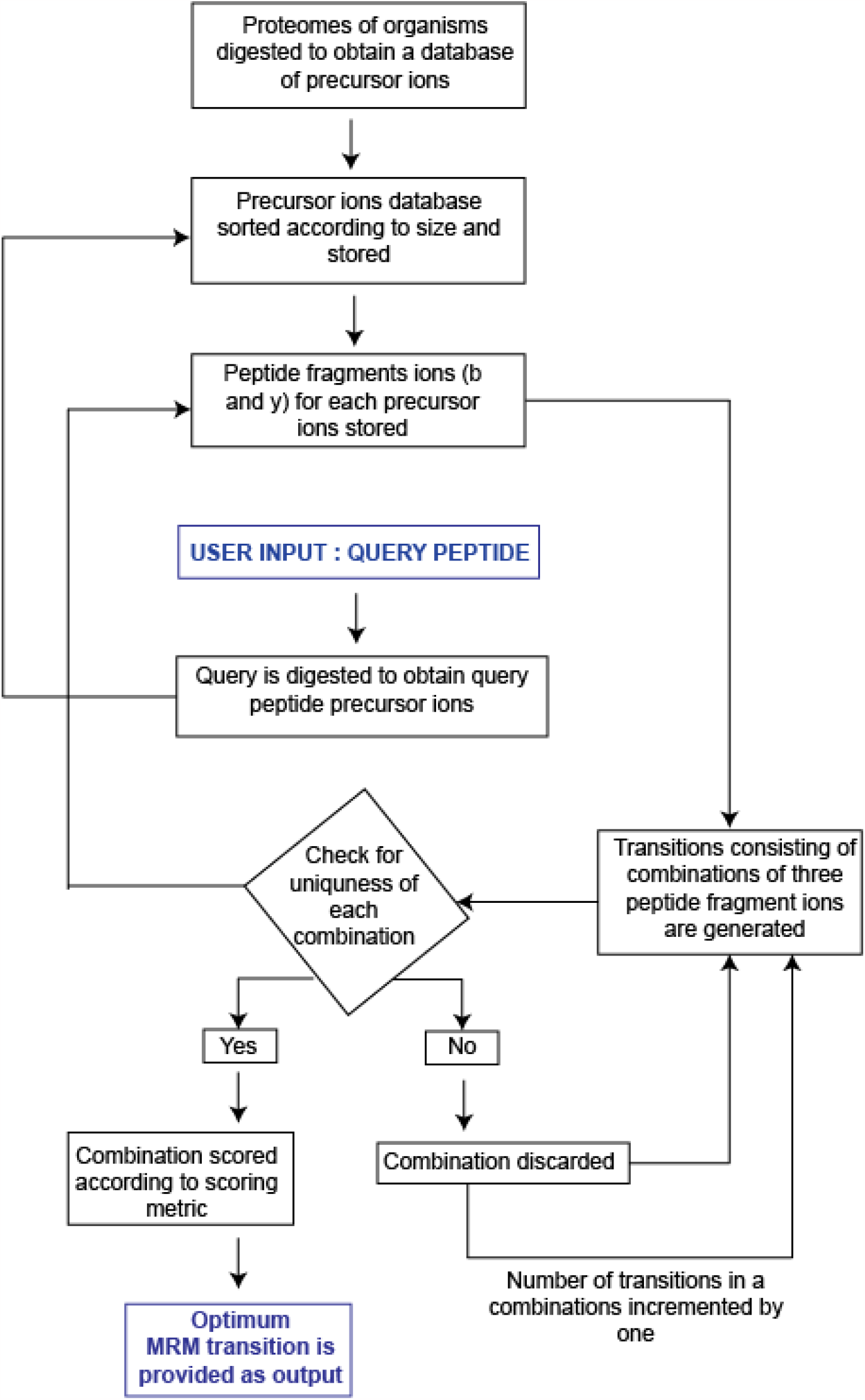
The algorithm for MRM design by DIGEST. The size of the peptide precursor and fragment ions is predetermined based on user input. If available, DIGEST provides more than one transitions as an output, with the transition scoring the highest ranked 1.

### Experimental validation of the design MRM module of DIGEST

We validated our MRM predictions experimentally by targeting an apolipoprotein panel of 12 proteins consisting of 20 peptides from pooled plasma samples of 10 healthy individuals. These proteins have high clinical value and are found to be biomarkers in several diseases ^18–20^. An accurate MRM design can therefore aid in rapid monitoring of several such biomarkers in a single run making it useful for medical applications. The three transition unique ions signature was obtained from DIGEST while five fragments per peptide were selected for MRM monitoring from SRMAtlas data. Of the 20 peptides, 18 apolipoprotein peptides could be identified with certainty using transitions obtained from DIGEST (**Table 2**). Therefore, our tool could design an MRM that could effectively detect most of the apolipoproteins. Interestingly, ‘Query Transition Values’ for the top three intense fragment ions of peptides for which Q3 transitions were selected from SRMAtlas compendium showed that most of these peptides did not make a unique ion signature (UIS). In other words, these top three intense fragments showed fragmentation redundancy with more than one peptide (**Figure 2**). Thus, indicating the requirement of a computational tool to aid in selection of fragment ion for UIS for any peptide.

**Table 2:** Experimental validation of UIS provided by DIGEST by comparing it with 6 most intense ions obtained from PeptideAtlas.

| Protein | Uniprot accession no. | Peptide sequence | Charge | PeptideAtlas (6 product ions) | DIGEST (UIS) |
| --- | --- | --- | --- | --- | --- |
| ApoA1 | P02647 | VSFLSALEEYTK | +2 | Yes | Yes |
|  |  | LLDNWDSVTSTFSK | +2 | Yes | Yes |
| ApoA2 | P02652 | EQLTPLIK | +2 | Yes | No |
|  |  | SPELQAEAK | +2 | Yes | Yes |
| ApoA4 | P06727 | SLAPYAQDTQEK | +2 | No | No |
|  |  | LAPLAEDVR | +2 | Yes | Yes |
| Apo(a) | P08519 | LFLEPTQADIALLK | +2 | No | No |
|  |  | TPAYYPNAGLIK | +2 | No | No |
| ApoB-100 | P04114 | ALVEQGFTVPEIK | +2 | No | No |
|  |  | FPEVDVLTK | +2 | Yes | Yes |
| ApoC1 | P02654 | EWFSETFQK | +2 | No (5) | Yes |
|  |  | TPDVSSALDK | +2 | Yes | - |
| ApoC2 | P02655 | ESLSSYWESAK | +2 | No (5) | Yes |
|  |  | TAAQNLYEK | +2 | No (5) | Yes |
| Apo C3 | P02656 | DALSSVQESQVAQQAR | +2 | Yes | Yes |
|  |  | GWVTDGFSSLK | +2 | Yes | Yes |
| ApoC4 | P55056 | DGWQWFWSPSTFR | +2 | No | No |
|  |  | AWFLESK | +2 | No | No |
| ApoD | P05090 | NILTSNNIDVK | +2 | Yes | Yes |
|  |  | VLNQELR | +2 | Yes | Yes |
| ApoE | P02649 | AATVGSLAGQPLQER | +2 | Yes | Yes |
|  |  | LGPLVEQGR | +2 | Yes | Yes |
| ApoF | Q13790 | SGVQQLIQYYQDQK | +2 | No | No |
| ApoL1 | O14791 | LNILNNNYK | +2 | No | No |
|  |  | VTEPISAESGEQVER | +2 | No | No |

**Figure 2:**
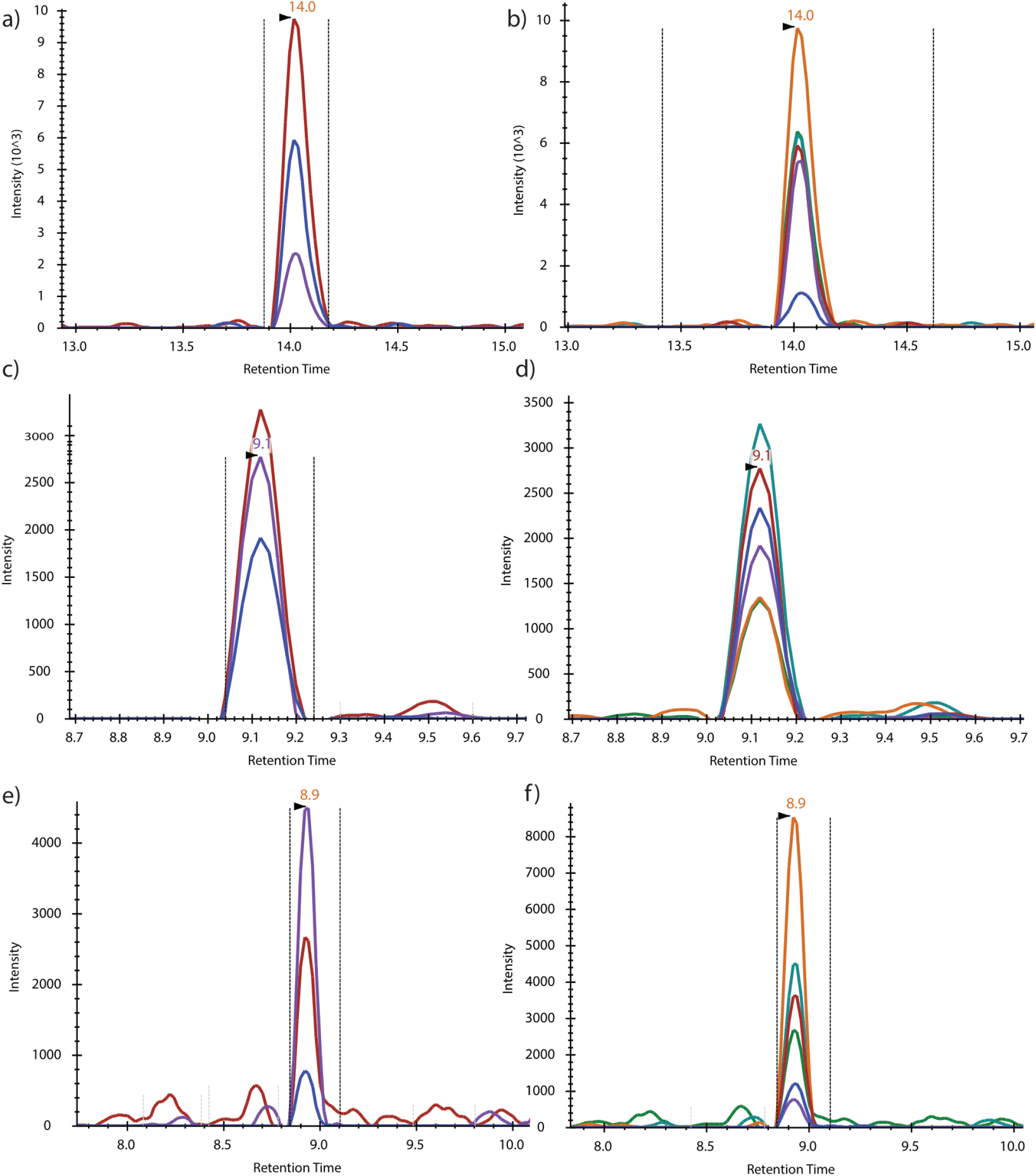
Extracted ion chromatograms depicting the intensity of signal observed for apolipoprotein sequences chosen for validation. Panel (a, c and e) depict extracted ion chromatogram for DIGEST ‘Design MRM’ UIS while panel (b, d and f) show extracted ion chromatogram of most intense fragment ions from PeptideAtlas datafile for peptides LLDNWDSVTSTFSK (+2), DALSSVQESQVAQQAR (+2) and LGPLVEQGR (+2) from Apo A1, Apo C3 and Apo E respectively.

### Other modules of DIGEST

#### a) Protein Digestion using various enzymes

In addition to MRM design, DIGEST enables users to digest a given peptide sequence using eight different enzymes to get digested peptides with either zero, one or two miscleavages using its ‘Protein Digestion’ module. Additionally, DIGEST can also be used to obtain all the precursor ions for the given peptide sequence allowing users to verify if there are other peptides in the proteome which may result in the same precursor ions. This may be done using its ‘Query by peptide sequence’ module. Similarly, the user may also provide transition values as precursor ion and product ion’s mass, charge and tolerance to obtain all the peptides in the proteome the input transition may be mapped to. This may be done using the ‘Query by transition values’ module. If the user chooses the result to include ‘All Q3-Ion Match’, only the peptides which generate all of the user provided Q3 *m/z* values are reported. Alternatively, to check for uniqueness of the signature, ‘Atleast one Q3-Ion Match’ may be selected to obtain any sequence that maps to even a single Q3 *m/z* value provided by the user.

#### b) Accounting for PTM modifications

Several proteins in the proteome undergo processing post-translation that leads to the covalent addition of chemical moieties. These post-translational modifications (PTM) of the proteins change their physical and functional properties, especially their mass^21^. To account for such changes in mass, DIGEST allows users to add 8 diverse and abundant modifications to amino acids while providing their query peptide sequence. The addition of a modified amino acid is facilitated by replacing the single letter amino acid code. This allows DIGEST to adjust the *m/z* values of the precursor and product-ions accordingly, increasing the accuracy of its output results.

## DISCUSSION

The success of an MRM/SRM assay in a complex matrix background is dependent on the choice of peptide sequence and the selection of the precursor and fragment ion combinations for the selected peptide. However, since peptides with different sequences can possibly share common transitions, the selection of these for an MRM assay remains challenging. Poorly designed assay may give ambiguous results due to interfering peptides that share transitions with the target peptide. We demonstrate that our tool can avoid these pitfalls by providing only those transitions [made of unique ion signature; UIS of three fragment ions (b and y) for a peptide] that uniquely map to the target peptide. DIGEST also allows users to query the uniqueness of a peptide sequence, and transitions based on user-determined mass tolerance in a user-defined proteome background.

Furthermore, we also demonstrate that a UIS of the order of three is sufficient to characterize∼98% of the proteome of organisms considered in this study (Humans, yeast, *E. coli, Arabidopsis thaliana* and SARS CoV2). DIGEST prioritizes the UIS for any query peptide sequence based on certain assumptions and ranks them based on their observability in experimental data. Thus, users can quickly determine the unique transition set for any peptide and get an experimentally observable output. ‘Query transition’ module can allow the users to perform UIS analysis on the data obtained in data dependent acquisition (DDA) mode and select experimentally observed transitions which are sufficient to uniquely assign peak group to a peptide.

Through experimental data we have shown that UIS prediction using ‘DesignMRM’ translates well to real world applications. We were able to identify almost all the Apolipoprotein peptides which were MS-observable by manually selecting 6 transitions per peptide from PeptideAtlas database. Using the minimal number of transitions (3 obtained from DIGEST) may also enable detection of a larger number of peptides in a single assay due to reduced scan cycle time. There have been several attempts to develop web-based tools to aid in MRM design^9,10^. However, most of these tools are currently unavailable, leaving a gap for the proteomics community to fill in. Future developments employing machine learning based approaches on existing proteomic experimental datasets to predict fragment ion intensity and digestion efficiency of a protein sequence with a selected protease may further optimize UIS output of MRM assays and provide more selective and sensitive transitions for MRM scans.

## ABBREVIATIONS

MRM: Multiple Reaction Monitoring
UIS: Unique Ion Signature
QTRAP: Quadrupole Ion Trap
TIQAM: Targeted Identification for Quantitative Analysis
MIDAS: MRM-Initiated Detection and Sequencing
ATQAS: Automated and Targeted analysis with Quantitative SRM

